# Scaffolding imagination: A role for medial frontal cortex in the expression of off-task thought

**DOI:** 10.1101/153973

**Authors:** Mladen Sormaz, Hao-ting Wang, Theodoros Karapanagiotidis, Charlotte Murphy, Mark Hymers, Daniel Margulies, Elizabeth Jefferies, Jonathan Smallwood

**Author notes:** Address for correspondence: Mladen Sormaz, Department of Psychology, University of York, Heslington, York, England.

## Abstract

We often think about people, places and events that are outside of our immediate environment. Although prior studies have explored how we can reduce the occurrence of these experiences, the neurocognitive process through which they are produced are less understood. The current study builds on developmental and evolutionary evidence that language helps organise and express our thoughts. Behaviorally, we found the occurrence of task unrelated thought (TUT) in easy situations was associated with thinking in words. Using experience sampling data, in combination with online measures of neural function, we established that activity in a region of anterior cingulate cortex / medial-prefrontal cortex (mPFC) tracked with changes in the expression of TUT. This region is at the intersection of two mPFC clusters identified through their association with variation in aspects of spontaneous thought: thinking in words (dorsal) and mental time travel (ventral). Finally, using meta-analytic decoding we confirmed the dorsal/ventral distinction within mPFC corresponding to a functional difference between domains linked to language and meaning and those linked to memory and scene construction. This evidence suggests a role for mPFC in the expression of TUT that may emerge from interactions with distributed neural signals reflecting processes such as language and memory.

## Main body

We spend upwards of one third of our daily lives engaged in thoughts and feelings related to events taking place in imagination rather than the here-and-now [1-3]. Although aspects of these experiences have beneficial associations with creativity [4], planning the future [5] and the reduction of psycho-social stress [6], their occurrence has also been linked to absent-minded error [7] and unhappiness [1, 2]. Consequently, the ability to organise self-generated thoughts is an important element of adaptive cognition [8].

Contemporary psychological theory has explored different mechanisms through which we influence spontaneous off task thoughts (for reviews see [9, 10]). One line of evidence suggests that mechanisms of executive control help supress task unrelated thoughts (TUT) allowing better focus on the task in hand when needed. This hypothesis is based on evidence that measures such as fluid intelligence [11] and attentional control [12, 13] are linked to reductions in the occurrence of TUT, especially in highly demanding externally-oriented tasks. Other studies have explored how we improve task focus through mental training or instruction [14-16].

Although these data explain how we maintain focus on external tasks, they do not make clear those processes that facilitate the expression of thoughts unrelated to events in the here and now [9, 10]. A comprehensive account of off task experiences requires an understanding of how they are generated, and how we shape this processes in line with our goals and desires [10]. Converging evidence from developmental [17, 18] and evolutionary literatures [19] emphasise the role that language processes play in shaping cognition, allowing it to explore realities other than those in the immediate environment [20, 21]. Based on this logic it is possible that we exploit language processes to facilitate the expression of TUT [22, 23] and in the current study we tested this hypothesis in two experiments that combined experience sampling with measures of neural function.

Contemporary neuroscientific theory, as well as meta-analytic evidence, highlight the default mode network (DMN) as important in the off task state [9, 24-26]. The DMN is a large-scale network anchored by hubs region in the anterior and posterior medial surface, as well as regions of lateral parietal cortex [27]. Online experience sampling using both functional magnetic resonance imaging (fMRI) and electroencephalography (EEG) have revealed activity in dorsal and ventral regions of anterior pre-frontal cortex / anterior cingulate cortex [28-30] as well as posterior cingulate cortex [31, 32] when participants are thinking about events rather than the here and now. These regions are also activated when participants explicitly engage in thoughts that mimic the experiences that participants produce in the off task state, such as thinking about the future or the past [33, 34]. Many of these regions of cortex also show a pattern of deactivation during demanding tasks (e.g. [27]) and these are also situations when off-task thought is more common [35]. As the DMN is thought to have an active involvement in mental content that can occur in off task thought, as well as exhibiting task induced deactivations, it is important to be able to differentiate neural signals associated with report of the off task state, from those whose behavior may simply the absence of conscious external task performance. In the current study we embedded thought probes into a cognitive task that alternated between a easier 0-back task and a more demanding 1-back task, allowing us to identify regions more sensitive to the expression of off task thought, rather than the demands placed by the external task.

The aim of the current project was to explore the neurocognitive processes that support the expression of off-task thought and to understand their relationship to language processes. We conducted two multi session experiments that combine experience sampling with measures of neural function, assessed using both task-based and task-free functional magnetic resonance imaging (fMRI). Experiment 1 used online experience sampling, in combination with task-based fMRI, to establish regions engaged by TUT. Experiment 2 measured intrinsic brain activity in a cohort of 153 participants using resting-state fMRI to characterise their functional organisation.

Using these data, the current study addressed several specific aims. First, we assessed the stability and reproducibility of the measures of experience sampling that form the basis of our study, as well as their consistency with prior investigations. Second, we demonstrate an association between the expression of task unrelated thought when task demands are low, and the tendency to think in words, providing support for the hypothesis that language processes can be engaged in the off task state. Third, using task based fMRI in combination with experience sampling, we identify a region of anterior cingulate / mPFC whose neural signal was sensitive to the expression of TUT. Fourth, we show that this region of mPFC falls at the intersection of clusters identified through their relationship to aspects of spontaneous thought: a dorsal cluster implicated in thinking in words and a ventral cluster implicated in the propensity for mental time travel. Finally, we performed a meta-analytic decoding to confirm the apparent dorsal/ventral distinction within the mPFC revealing a functional division between language (dorsal) and memory (ventral). Together these results underline the importance of the mPFC in supporting the expression of mental content in the off-task state.

## Results

### Behavioural analysis

In both Experiments we measured the contents of experience in a task context that alternates between a demanding condition, that places an external demand on working memory (1-Back), and a less demanding condition, that does not (0-Back). The latter has lower task demands and so provides greater opportunity for TUT and its associated neural processing [36] (Fig 1 A). Consistent with expectations and prior studies [36], participants performed less efficiently on the more difficult task in both Experiments (Scanner: 1-Back, M = .116 (SE = .001), 0-back, M = .12 (SE =.002) t (28 = 2.43, p<.05; Lab: 1-Back M =. 097 (SE =.003), 0-Back M =.124 (SE =.003) t (151) = 17.5, p<.001).

**Fig 1.**
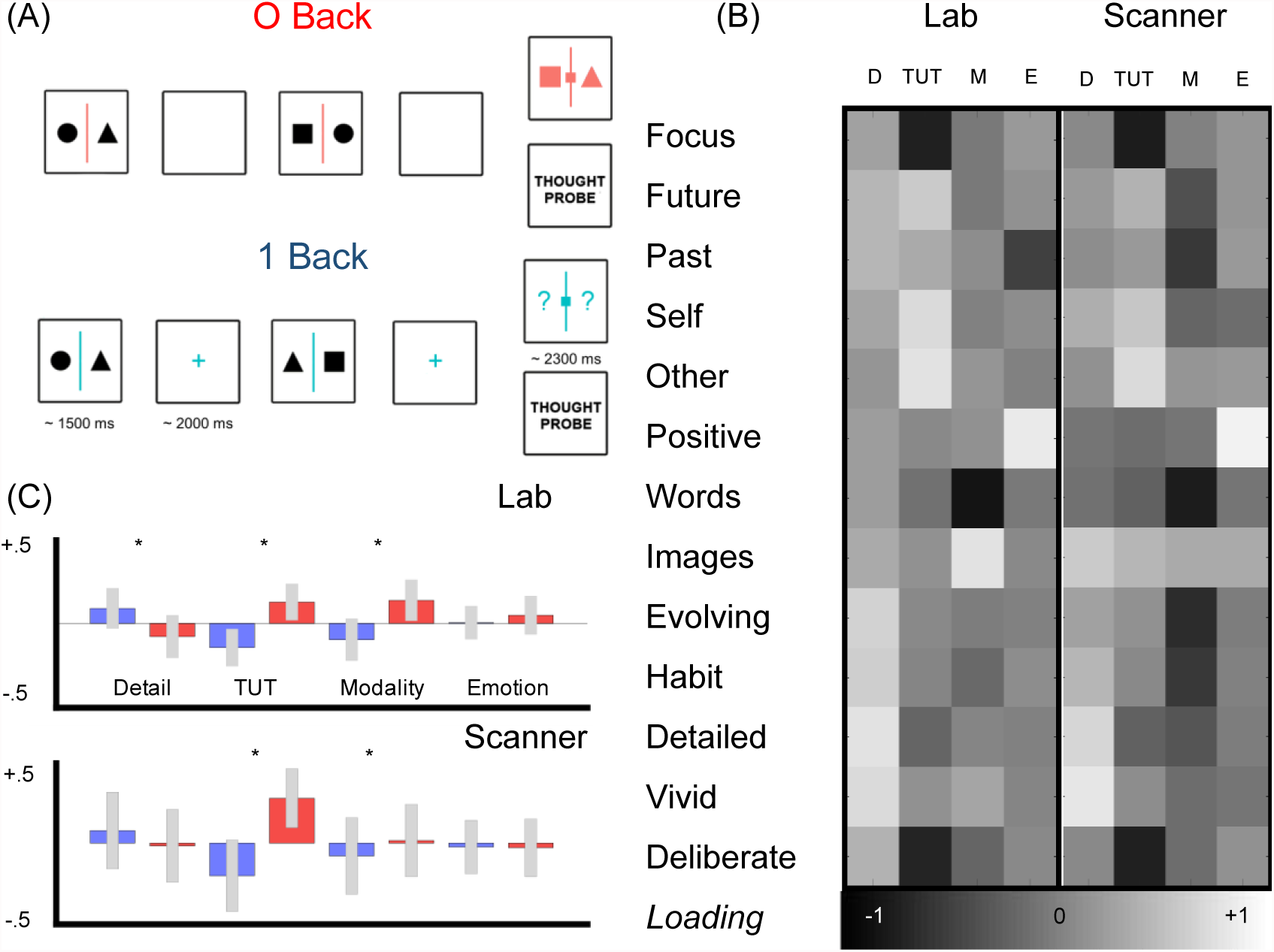
Assessing the contents of experience. (A) In both Experiment 1 and 2 we measured experience while participants performed a task in which they viewed pairs of shapes (triangles, squares and circles) and made intermittent decisions about the spatial location of these shapes at catch trials. The task was organized into alternating blocks that were either easy, because the decision was made in the context of the available evidence (0 Back) or more difficult because the participants decisions depended on information present on the prior trial (1-back) requiring them to maintain a task relevant stimulus representation throughout the block. (B) At irregular intervals we collected self-reports throughout this task using Multi-dimensional Experience Sampling (MDES) which we decomposed using exploratory factor analysis revealing four dimensions: Detailed and vivid experiences (D), Task Unrelated Thought (TUT) about people and other times, Images or Words (Modality, M) and Emotion (E). (C) Projecting these dimensions back onto the tasks revealed that in the more difficult task TUT was lower, and thoughts more often took the form of Words.

**Fig 2.**
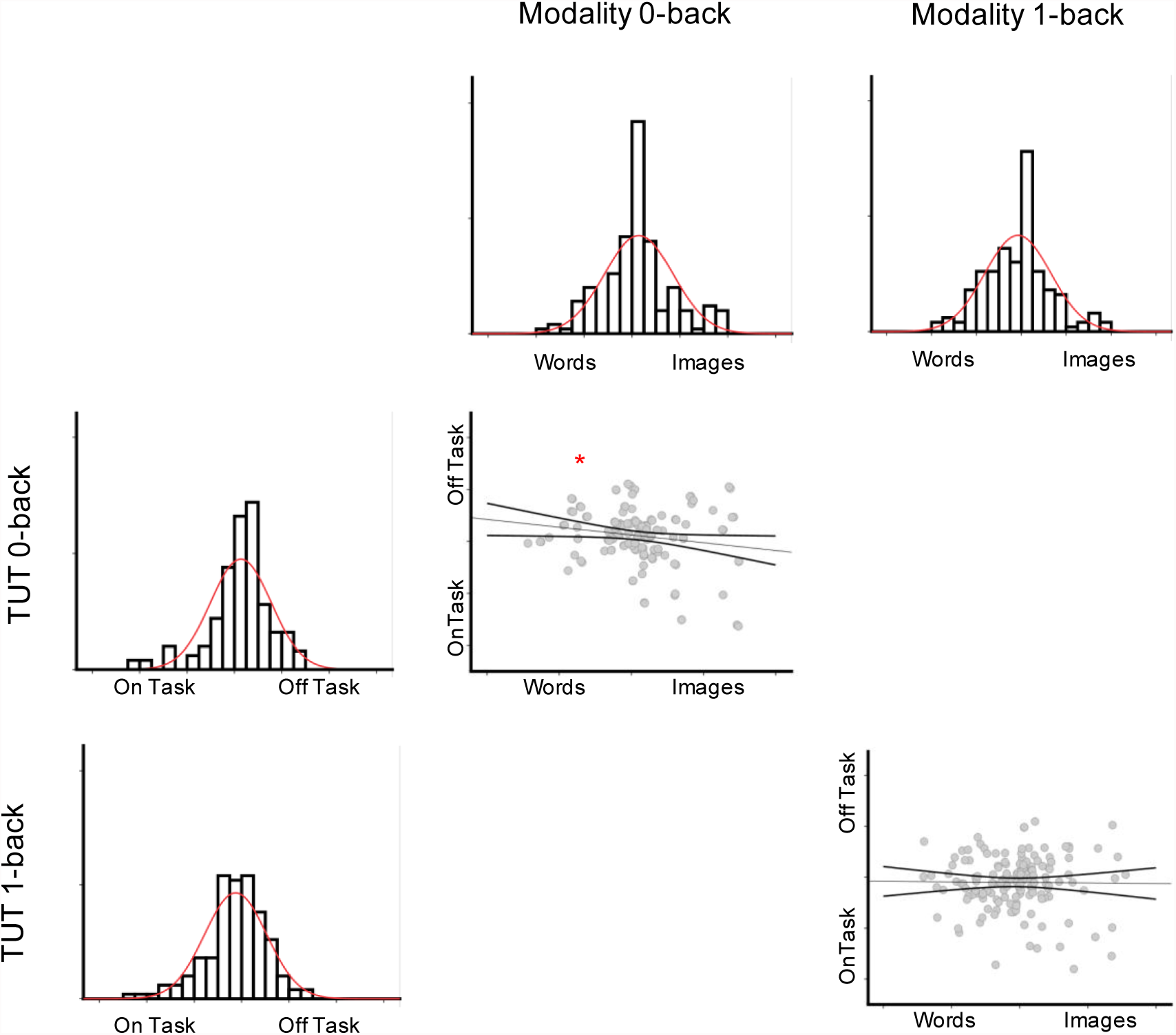
Association between task unrelated thinking and the modality of thought in easy and hard tasks.

We measured experience using multi-dimensional experience sampling (M-DES), a technique in which participants are periodically interrupted while they perform a task and asked a series of questions regarding their experience at that moment in time [37, 38]. As in our prior work we decomposed the self-reported data generated by MDES using exploratory factor analysis at the trial level, to reveal their underlying dimensional structure. With respect to the current data, this reveals dimensions describing the level of vividness and detail in ongoing experience (*Detailed*, vivid experiences, D), the relationship of the experience to the ongoing task (*Task-unrelated Thought*, TUT, typically social thoughts about the future), whether the experience is in the form of images of words (*Modality*, M), and their emotional status (*Emotion*, E). The result of this decomposition procedure is presented in Fig. 1B in the form of a heat map.

These patterns are broadly consistent with the patterns seen in our prior studies [37, 38] and importantly we see qualitatively similar patterns inside and outside of the scanner. Quantitative comparisons revealed that components from each context were correlated (See Table S1) allowing us to generalize across experimental contexts. In addition, in both the scanner and the laboratory, thoughts in the more difficult task were more deliberately focused on the task (*Lab*, *t* (152) = 4.64, *p* < .001; *Scanner*, *t* (29) = 5.89, *p* < .001) and rated as more often in the form of words than images (*Lab*, *t* (152) = 4.06, *p* < .001; *Scanner*, *t* (29) = 2.24, *p* < .05) (see Fig 1 C). Together this analysis demonstrates correspondence between experience as described in the scanner and in the laboratory. It also shows that in both contexts the manipulation of task difficulty systematically changes the participants experience, leading to greater TUT in the easier task 0-back task and a greater tendency to think more in words in the harder 1-back task.

Our next analysis replicated the relationship between fluid intelligence and reduced TUT in more demanding task conditions seen in prior studies [3, 7, 11]. We analyzed the loading of the component associated with TUT using a mixed Analysis of variance (ANOVA). This had a single within-participants factor with repeated measures on each task. In this analysis we included the average score on Raven’s Progressive Matrices as a continuous between participant factor. This revealed a Task X Fluid intelligence interaction [F (1, 151) = 4.5, p<.05] indicating the relationship between task focus and fluid intelligence varied significantly across tasks. Follow-up correlations indicated a negative relationship in the 1- Back task (r (152) = -.17, p < .05) but no relationship in the 0-back task (r (152) = .01, p = .88). This analysis replicates prior work that shows that executive control can suppress TUT in difficult task.

Our final behavioural analysis explored whether there was any association between the expression of TUT and the tendency to think in words. We analyzed the loading of the component associated with Modality using a mixed Analysis of variance (ANOVA). This had a single within-participants factor with repeated measures on each task. In this analysis we included the scores describing the propensity for TUT in the 0-back and the 1-back tasks as separate predictors. We also included the average score on Raven’s Progressive Matrices as a control. This analysis revealed a Task X TUT O-back [F (1, 149) = 4.095, p < .05] interaction. This interaction is attributable to the fact that in the easy 0-back task, there was a weak yet significant correlation between greater TUT and reports of thinking in words (r = -.167, p <. 05, see Figure Two, upper panel). There was no association between the modality of thought and the level of task focus in the more demanding 1-back task. This pattern suggests that under conditions when task demands are reduced off-task thought tends to be expressed in words.

These behavioural analyses demonstrate three aspects of the generalizability of our data: (i) a correspondence between our scanning and laboratory based measures of experience, allowing us to generalize across contexts, (ii) shows that task difficulty manipulation changes experience in terms of it’s level of task focus and the modality of experience, demonstrating our task manipulation impacts upon experience and (iii) shows our paradigm is sensitive to measures of fluid intelligence in a manner that is consistent with prior studies. They also provide basic support for the hypothesis that language processes are important in off-task thought since under non-demanding task conditions the expression of task unrelated thought tends to be associated with thinking in words.

## Functional magnetic resonance imaging

### Identifying neural regions associated with task unrelated thought

Our first aim with the task-based imaging was to determine the neural regions that contains signals sensitive to the expression of off task thought. Using the task-unrelated dimension generated by MDES as a regressor of interest in the online fMRI data collected in Experiment 1 (see Methods), we identified a region of mPFC associated with the occurrence of TUT (shown in green in Figure 3). This region has been observed in prior studies exploring off task thought (e.g. [28, 29, 31, 39]). Our analysis suggests that the neural signal in this region of mPFC is more sensitive to the expression of TUT than it is to the deactivations associated with high levels of task demands. To understand this region fully we performed a seed based functional connectivity analysis using this region as a mask (see Methods). This revealed a pattern of functional connectivity that showed strong connectivity with the hubs of the DMN (anterior mPFC, posterior cingulate cortex and bi-lateral angular gyrus). In addition this analysis highlighted the anterior insula, thalamus and the striatum. This spatial pattern corresponds to the combination of two-large scale networks – the DMN and the cingulo-opercular or saliency network [40, 41].

**Fig 3.**
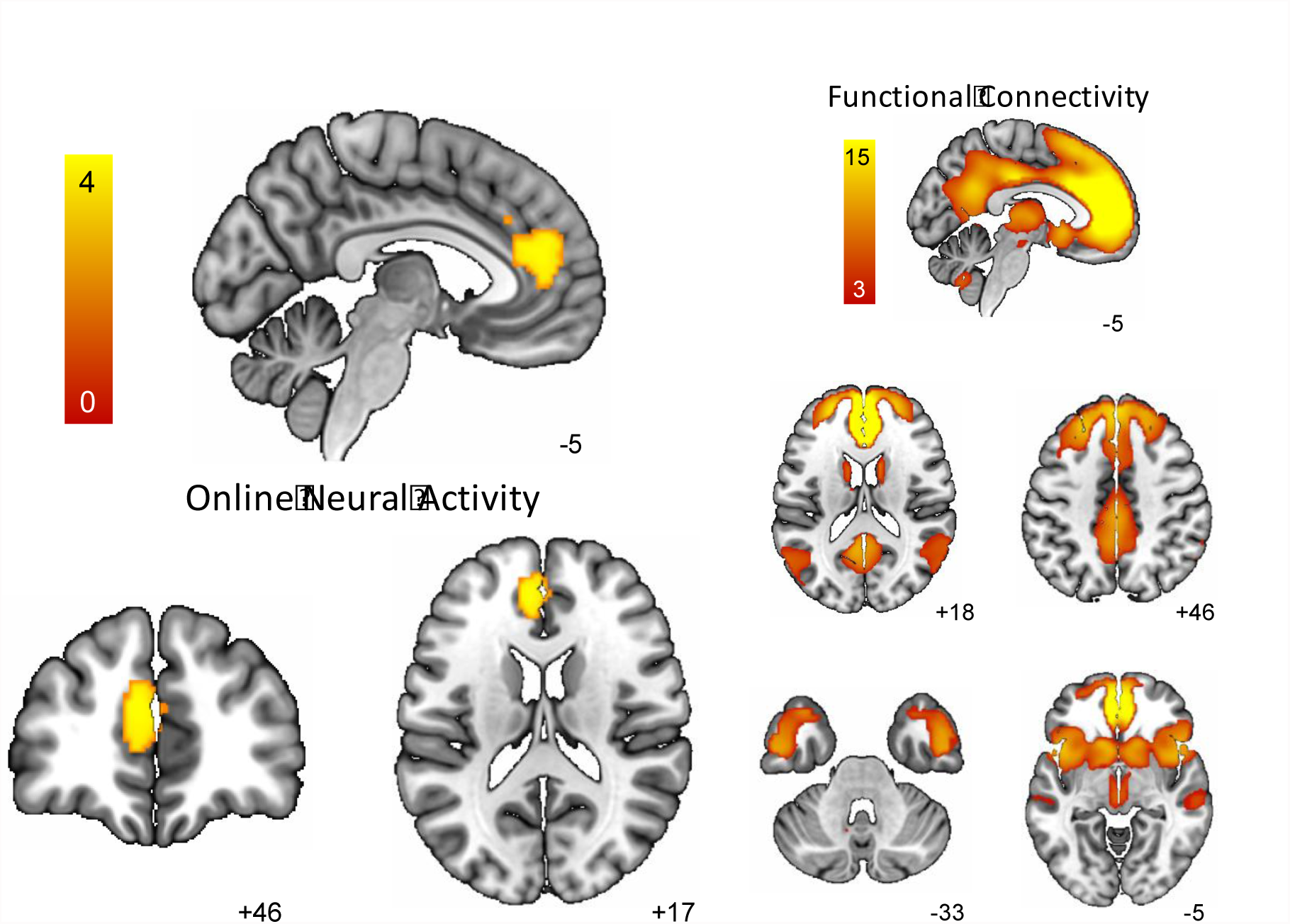
Regions whose activity is associated with ongoing task unrelated thought (TUT). All brain images were thresholded at Z = 3.1 and corrected for multiple comparisons at p < .05 FWE.

Next we identified those regions that showed a response profile that varied with level of external demands. Regions of cortex showing evidence of task-induced deactivation are shown in red in Figure 4. These regions included regions of medial prefrontal and posterior cingulate cortex, rostro-lateral pre-frontal cortex and regions of the temporal lobe. Many of these regions fall within the default mode network [24]. It can be seen that the regions of task-induced deactivation were adjacent to, and largely non-overlapping with the mPFC region sensitive to the expression of TUT. We also identified regions that show a stronger response when external task demands are increased. Regions more active in the more demanding 1–Back task are shown in blue and include dorso-lateral prefrontal cortex, pre-supplementary motor cortex, posterior middle temporal gyrus and lateral parietal regions, including the inter-parietal sulcus. Many of these regions fall in the fronto-parietal or multiple demand network [42] that are linked to task related cognition when external task difficulty is increased [43].

**Fig 4.**
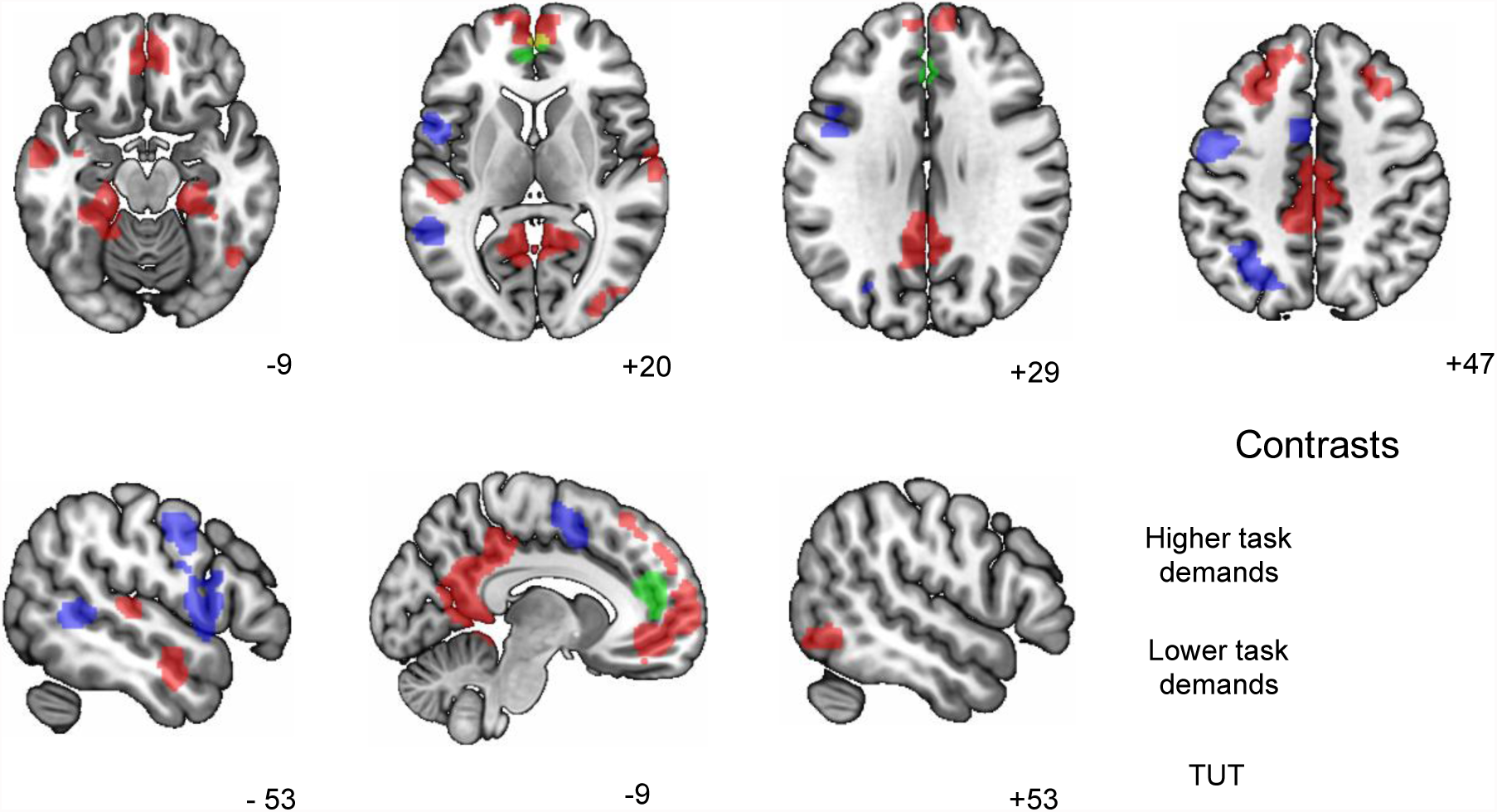
Localising regions associated with high (blue) and low (red) task demands. Regions that reflect the ongoing experience of TUT and shown in green. All brain images were thresholded at Z = 3.1 and corrected for multiple comparisons at p < .05 FWE.

### Identifying the intrinsic architecture supporting different features of experience

Having established a region of mPFC that shows sensitivity to the expression of TUT, as well as those regions that respond to the demands of the task, we explored the intrinsic architecture associated with these spatial maps and examined whether they are sensitive to individual differences in types of experiences. Our first analysis used the mPFC region, identified as important for the expression of TUT, as a seed in a functional connectivity analysis (see Methods). This revealed a pattern of functional connectivity with a region of motor cortex that varied differentially with the modality of cognition in each task (Figure 5, Left hand panel). It can be seen in the scatter plots that this region was more coupled for individuals whose experience was in the form of words than images in the easy 0-back task, and more decoupled for individuals who thought in words in the harder 1 back task. This reflects a pattern of dMPFC-motor coupling that varied with the extent of task differences in the modality of experience.

**Fig 5.**
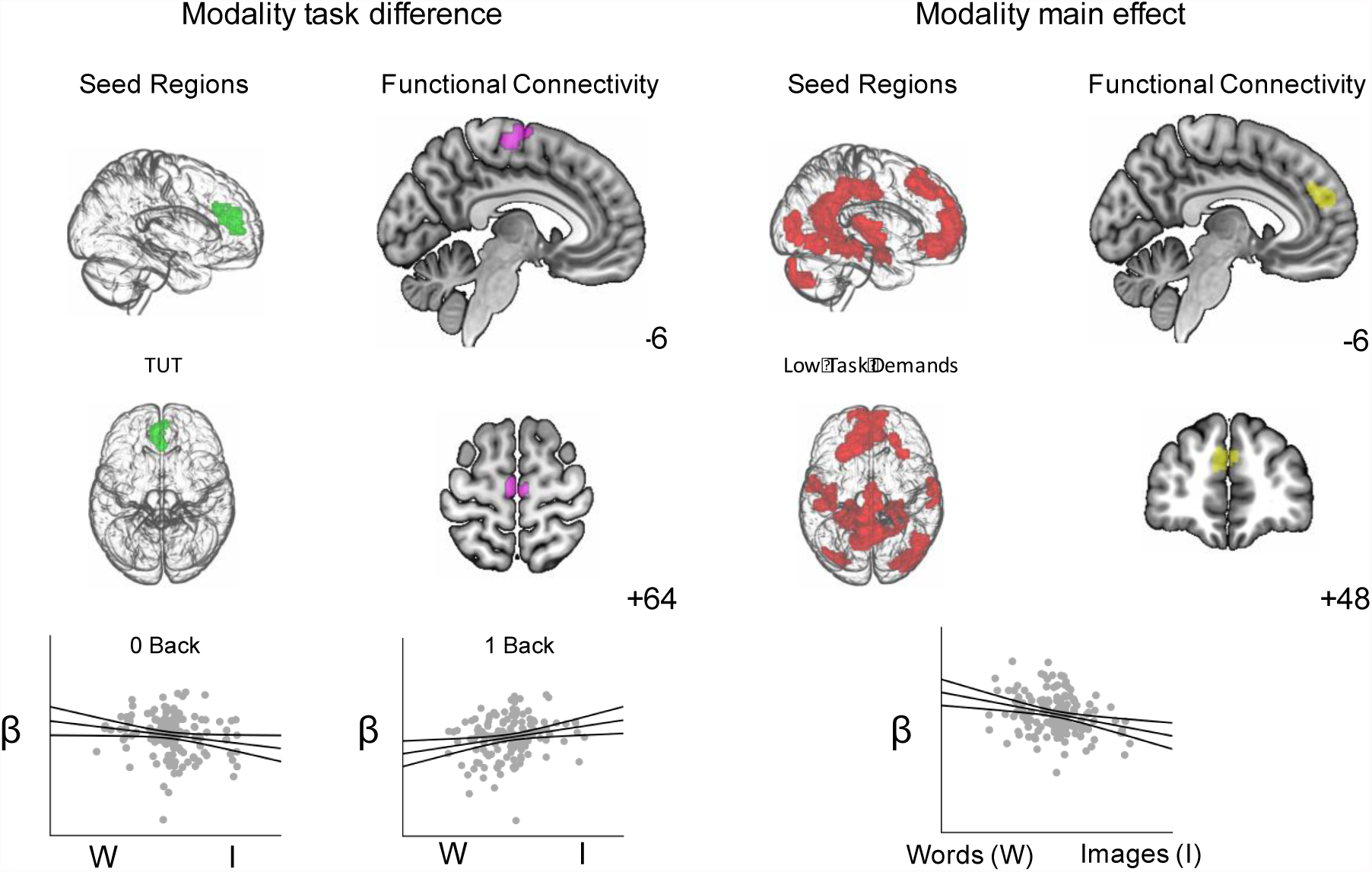
Intrinsic architecture supporting variations in the modality of thought. *Left hand panel.* A region of dorsal mPFC linked to TUT (green) had increased connectivity with a region of motor cortex (purple) for individuals who thought more in words in the easier 0-back task and images in the harder 1-back task. *Right hand panel.* Regions showing more activity in the easy 0 back task (red) had increased connectivity with the dorsal mPFC (yellow) for individuals who describe their experiences as consisting of words. The seed regions in these analyses are masks generated by the contrasts presented in Figure 4. All brain images were thresholded at Z = 3.1 and corrected for multiple comparisons at p<.05.

Our next analysis used the network of regions identified as active in the non-demanding task from Experiment 1 as seed regions. This revealed two significant results: (i) more positive experiences were associated with increased communication with a region in right lingual gyrus (see SF2) and (ii) experiences that took the form of words, rather than images, were associated with greater connectivity with a region of dorsal mPFC (see Fig 5 Right hand panel). This latter result suggests that the tendency to think in words is linked to a pattern of functional connectivity with the dorsal mPFC from regions of cortex showing greater activity when external task demands are low.

In combination these analyses highlight the broader mPFC region as important for the intersection between TUT and language processes. Fig 6 (left hand panel) displays the spatial overlap of these two analyses. The online experience sampling analysis (presented in green) is adjacent and semi-overlapping with the dorsal mPFC region linked to the tendency to think in words (presented in red). In addition, our prior work established a more ventral region of mPFC, which was coupled to the hippocampus and linked to episodic aspects of mind-wandering [37] (a pattern which we replicated in Experiment 2, see Fig S2). This cluster is presented in blue. It falls largely within ventral aspects of the mPFC region active during off-task thought. It is apparent from this figure that the region of mPFC active during off-task thought falls at the intersection of both dorsal and ventral cluster related to language related processing and memory related processing aspects of experience respectively.

**Fig 6.**
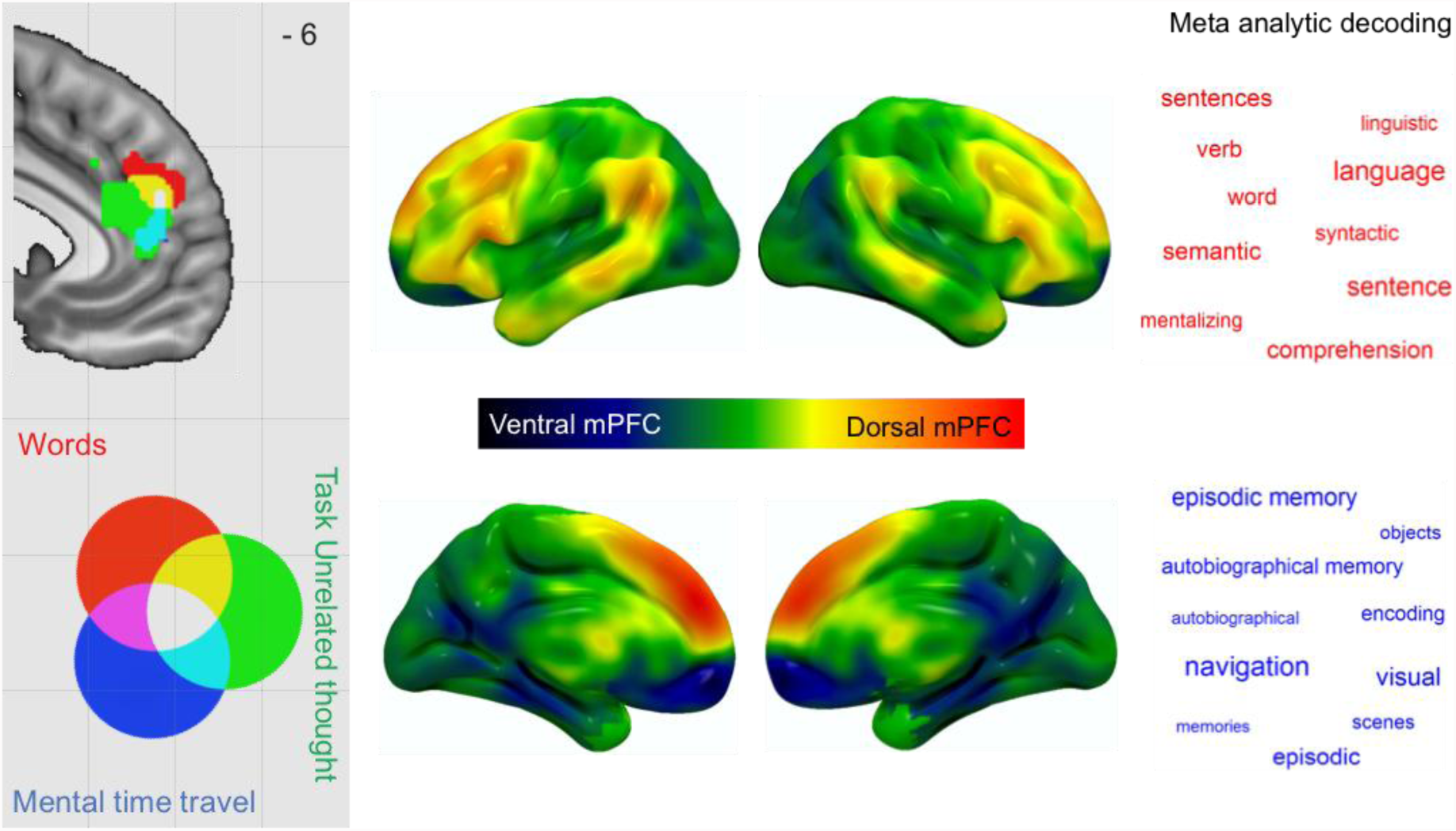
Right hand panel. A region of mPFC recruited during TUT (indicated in Green) overlaps with both the cluster linked to thinking in words from the current study (indicated in red) and a ventral region linked to episodic memory in a prior study by Karapanagiotidis and colleagues (indicated in blue). *Middle panel.* Differential connectivity in an independent set of 140 participants revealing regions showing greater connection with the dorsal mPFC region (coloured red-yellow) and ventral mPFC regions (coloured in blue-green). This map is unthresholded (see Fig S4 for the thresholded equivalent). *Left hand panel.* Meta analytic decoding using Neurosynth revealed functional associations with memory for regions with stronger connectivity to ventral mPFC (indicated in blue) and those with stronger connectivity to dorsal mPFC (indicated in red).

### Meta analytic decoding

Our final analysis sought to understand the functional significance of this spatial distribution of activity within the mPFC. We contrasted connectivity from the ventral and dorsal regions of the mPFC presented in Fig 6 in an independent set of 140 participants from a publicly available repository of resting state fMRI (see Methods). This revealed stronger functional connectivity from the dorsal mPFC cluster to regions of lateral pre-frontal, parietal and temporal regions (see Fig 6 Middle panel, see also supplemental Fig 4). In contrast, stronger connectivity was observed from the ventral mPFC to the posterior cingulate as well as to medial regions of the temporal lobe. It is noteworthy that the difference between these two regions corresponds broadly to the dorso-medial and medial-temporal subsystems of the default mode network identified by Andrews-Hanna and colleagues [24, 44]. To understand the functional significance of these difference in connectivity, we performed a meta-analysis of this differential spatial map using Neurosynth [45]. Connectivity that was greater for dorsal rather than ventral mPFC was associated with terms such as “language”, “sentences” and “semantic”. Regions showing greater connectivity with ventral than dorsal mPFC was linked to terms such as “episodic memory”, “autobiographical memory” and “scenes”. This provides corroborative meta-analytic evidence that links the regions of mPFC, identified through its relationship to experiences, is activated by functional studies using task paradigms that probe the appropriate domain.

## Discussion

Our study highlights an area of anterior cingulate cortex and mPFC as playing an important role during off-task thought. Using experience sampling with online fMRI, we were able to demonstrate that the signal in the aCC / mPFC tracks with the expression of TUT rather than an absence of external task demands. This region has been observed in prior studies of TUT [28, 31, 32]. Functionally this region shows connections to both the DMN, as well as to so-called saliency or cingulo-opercular system [40, 41]. In the context of contemporary accounts of spontaneous thought, interactions between the DMN and the cingular-opecular network are thought to reflect the increased salience associated with internal processing during states such as task-unrelated thought [9].

Our study raises the possibility that the mPFC is important in off-task thought because it can integrate different neuro-cognitive processes that are distributed across the cortex, and reflect higher forms of cognition, such as memory or language. Behaviourally we found that when task demands were minimal, task unrelated thinking was associated with experiences characterised as words. Our analysis combining online experience sampling with fMRI allowed us to establish a region of mPFC / aCC that was located at the intersection of two regions identified through an analysis of the intrinsic architecture linked to spontaneous thought: a dorsal region linked to the tendency to think in words, and a more ventral region linked to episodic aspects of ongoing cognition. Meta-analytic decoding linked the connectivity of this dorsal region to functions such as “sentences”, “comprehension” and “language, while the ventral region of mPFC was linked with terms such as “episodic memory”, “autobiographical memory” and “navigation”. Together these analyses support an integrative role of the mPFC during off task thought since its functional and spatial profile would allow the integration of information from large-scale systems that support different aspects of higher order cognition, such as memory and language.

More speculative support for the view that mPFC plays a role in off-task thought through a process of integration comes from our analysis of how the intrinsic organization of this region relates to spontaneous thought. We observed a pattern of coupling with a region of primary motor cortex that predicted the modality of a participant’s experience in a task specific manner, a dissociation that may reflect the joint role that the motor system plays in action and imagination [46-48]. In the more demanding 1-back task, where performance depends upon rehearsing visual spatial features encoded into memory, thinking in words is associated with mPFC-motor cortex decoupling. In contrast, in the less demanding 0-back task, where off-task experiences are more often in the form of words, the coupling of the mPFC with motor cortex may reflect the integration of an embodied motor contribution to cognition, perhaps providing supporting imagined actions [49]. Consistent with this latter possibility we have previously found that hippocampal connectivity with a similar region of motor cortex is linked to the process by which individuals generate more specific personal goals in imagination [50]. Note however that this interpretation should be treated as speculative in lieu of more direct evidence of the functional significance of interactions between motor and pre-frontal cortex and off task thought.

Finally, our study has important implications for psychological accounts of how we organise thoughts that are independent of the external environment. We replicated prior studies that found measures such as fluid intelligence act to control the occurrence of TUT when task demands are high [7, 11]. This process of executive control helps regulate the context in which off-task thoughts emerge and so limits the cost of TUT states on ongoing performance [8, 12]. In contrast, our study suggests that a greater expression of TUT when the environment lacks notable external demands is linked to a bias to thinking in words. This suggests that we may capitalise on language processing to facilitate the expression of our thoughts in the off task state [10]. In this way our data supports perspectives from both developmental psychology [18] and cognitive psychology [51] which suggest inner speech provides scaffolding that help us pursue trains of thought concerning times and places other than where we are now. More generally, if this account is correct it suggests that although both fluid intelligence and language influence the off-task state, they do so in different ways. Fluid intelligence reduces the occurrence of TUT when external demands are high, improving focus on complex external tasks. In contrast, language processing facilitates the expression of TUT when the cognitive demands of the external environment are minimal. The contrasting role of fluid intelligence and language may reflect the dissociation between process and occurrence that has been argued to be important in understanding off task thought [10].

Before closing it is worth considering the limitations of our findings. Although our data suggest processes such as language may be important role in the expression of task unrelated thought, there are different mechanisms that this association could reflect. For example, language could help organise off-task thought by providing a grammatical, or syntactical structure, that helps order cognition over relatively short time scales (e.g. [52]). Thinking in words may also provide the conceptual structure that underpins off-task thinking, by providing access to representations that makes up our semantic knowledge of the world [53, 54]. Both of these possibilities are consistent with our meta-analytic evidence: Decoding the connectivity of the dMPFC emphasised both functions linked to linguistics (e.g. “syntactic”, “verb”) and meaning (e.g. “semantic”, “comprehension”). It will be important for future research to explore how components such the meaning and phonology of language processing shape the expression of off task thought.

Our study also leaves open whether the contribution of language processing, or memory, allows off task thoughts to function in a more adaptive manner. For example, thinking in words has been shown to be linked to the capacity to introspect [22] and maybe important in planning the future [23]. When tasks demands are low, the expression of TUT is often linked to more positive outcomes, such as a more patient style of decision-making [55, 56], or better working memory [13, 57]. Since our study links thinking in words, to the expression of TUT in easy task contexts, it is possible that the contribution of language to off-task thought facilitates positive aspects of this state, such as the process of autobiographical planning [5]. Alternatively, the contribution of language process during the off-task thought could explain why this experience can be detrimental to on-going reading comprehension [58]. Similarly, verbal worry can also be problematic in states of heightened anxiety, by acting to lengthen the duration of an episode [59]. Future work should aim to understand whether the differential contribution of language and memory to the off task state determines particular functional outcomes linked to the mind-wandering state and does so in a manner that is generally beneficial or detrimental to an individuals well being.

## Methods

### Participants

All volunteers provided informed written consent and were paid either £20 or given course credit for their participation. They were right-handed, native English speakers, with normal/corrected vision and no history of psychiatric or neurological illness. Note that participants in Experiment 1 also took part in Experiment 2 and that Experiment 2 was conducted first. Both studies were approved by the University of York Neuroimaging Centre ethics committee.

#### Experiment 1

Thirty-four participants (15 females; mean ±SD age = 22 ± 2.2 years) were recruited for experiment 1. Four participants were excluded at the data analysis stage due to extreme motion in the FMRI scanner in more than 50% of runs.

#### Experiment 2

One hundred and sixty four participants were recruited for whom 153 completed the full testing sessions and were part of the data analysis cohort (95 females; mean ±SD age = 20.1 ±2.1 years). They were recruited within the same exclusion criteria and ethical guidelines as those in Experiment 1.

#### Independent Sample

We also used an independent dataset to provide independent confirmation of functional connectivity results. These data were obtained from a publicly available dataset: the Nathan

Kline Institute (NKI)/Rockland Enhanced Sample and contained 141 subjects. Full details of this sample can be found in [31].

### Procedure

#### Experiment 1

Participants completed two one-hour long fMRI scanning sessions completed on separate days, at least 24 hours apart. Each session consisted of four 9 minute runs that alternated between the 0-back and 1-back conditions (see Figure 1).

#### Experiment 2

Participants in experiment 2 underwent the same fMRI resting state scanning and behavioural testing procedures as outlined in [60].

### Procedure

#### Task paradigm

Non-target trials in both 0 back and 1-back conditions were identical, consisting of black shapes (circles, squares or triangles) separated by a coloured line signifying whether the condition was 0 back or 1-back (mean presentation duration = 1050 ms, 200 ms jitter). The non-target trials were followed by presentation of a black fixation cross (mean presentation duration = 1530 ms, 130 ms jitter). Non targets were presented in runs of between 2 and 8 with a mean of 5 following which a target trial or a MDES probe was presented. In neither condition do participants make a behavioural response to the non-target trials.

The target trials differed by condition. In the 0-back condition, the target trial was a pair of coloured shapes presented either side of a coloured line with a probe shape in the centre of the screen at the top. Participants had to press a button to indicate whether the central shape matched the shape on the left or right hand side of the screen. In the 1-back condition the target trial consisted of a coloured question mars presented either side of a coloured line with a probe shape in the centre of the screen. Participants had to indicate with a button press a button whether the central shape matched either the shape on the left or right side of the screen on the previous (non-target) trial.

### Experiential Assessment

In both experiments the content of thought was measured using Multidimensional experience sampling (MDES). This involved the presentation of 13 questions presented in Supplementary Table 2.

### Ravens advanced progressive matrices

The Ravens Advanced Progressive Matrices (RAPM; [32]) measured ‘fluid intelligence – that is the ability to make sense and meaning out of complex non-verbal stimuli. In order to complete the task participants were tasked with finding new patterns and relationships between the stimuli. The RAPM used in the current study contained two tests: (i) practice test - containing 2 problems and (ii) the full test – containing 36 problems. For each problem a set of 9 boxes (ordered in a 3x3 design) were shown on the screen. All but one box contained a pattern. At the bottom of the screen were 4 additional boxes, each containing a unique pattern. Participants were required to select out of these 4 potential boxes which pattern should go in the empty box. During the practice phase participants were given online feedback outlining whether their response was correct and, if not, how they should decide which box was the correct answer. If participants had any further questions, then they were instructed to ask the experimenter before starting the main experiment. During the full test no feedback was given. Participants were given 20 minutes to complete as many problems as they could, the problems got progressively more difficult.

#### Experiment 1

MDES probes occurred at 6 points during each run participants completed 13 questions about their experience. Responses were made on a 4 point Likert scale. Participants had 5 seconds to respond to each question and in total the probe question points were of a fixed 60 second duration. In each run there were an average of 3 question periods in the 0 back condition and in the 1-back condition.

#### Experiment 2

In Experiment 2 MDES probes occurred on a quasi-random basis to minimize the likelihood that participants could anticipate the occurrence of a probe. At the moment of target presentation that there was a 20% chance of a MDES probe instead of a target with a maximum of one probe per condition block of 0-back and 1-back. In each session, an average of 14 (*SD* = 3.30, range 6 – 25) MDES probes occurred; in the 0-back condition an average of 7 (*SD* = 2.36, range 2 – 14) MDES probes occurred and in the 1-back condition an average of 7 (*SD* = 2.24, range 1 –15) occurred.

#### MRI Image acquisition

##### MRI acquisition

MRI functional and structural parameters for the task based and resting state fMRI scans were identical. Both structural and functional data were acquired using a 3T GE HDx Excite MRI scanner utilising an eight-channel phased array head coil (GE) tuned to 127.4 MHz, at the York Neuroimaging Centre, University of York. Structural MRI acquisition in all participants was based on a T1-weighted 3D fast spoiled gradient echo sequence (TR = 7.8 s, TE = minimum full, flip angle= 20°, matrix size = 256 x 256, 176 slices, voxel size = 1.13 x 1.13 x 1 mm). Resting-state activity was recorded from the whole brain using single-shot 2D gradient-echo-planar imaging (TR = 3 s, TE = minimum full, flip angle = 90°, matrix size = 64 x 64, 60 slices, voxel size = 3 x 3 x 3 mm^3^, 180 volumes). A FLAIR scan with the same orientation as the functional scans was collected to improve co-registration between subject-specific structural and functional scans. MRI acquisition details of the independent sample can be found in [31].

#### Data Analysis

##### 0 back and 1-back task performance

In both experiment 1 and 2 we recorded the mean accuracy and reaction time (RT) for participants in the 0 back and 1-back experimental condition. From this we calculated an efficiency score (percent accuracy correct/ RT in milliseconds) based on the inverse efficiency score described by [33] although we used percent correct as the numerator instead of percent incorrect due to the low error rate across participants, we then z-scored this efficiency score to the mean for each participant.

##### Multi-Dimensional Experience Sampling (MDES)

In both experiments 1 and 2 we used principal components analysis (PCA) with varimax rotation in SPSS version 24 to decompose the dimensionality of both the experience-sampling data. We used the outcome of scree plots (see supplementary figure 3) from the PCA analyses carried out in experiments 1 and 2 to determine that 4 principal components should be extracted. These four principal components were then used to model fMRI data in experiments 1 and 2. See results section for the description of principal components extracted.

##### Task based and resting state fMRI

In both Experiment 1 and 2 functional and structural data were pre-processed and analysed using FMRIB’s Software Library (FSL version 4.1, http://fsl.fmrib.ox.ac.uk/fsl/fslwiki/FEAT/). Individual FLAIR and T1 weighted structural brain images were extracted using BET (Brain Extraction Tool). Structural images were linearly registered to the MNI-152 template using FMRIB’s Linear Image Registration Tool (FLIRT). The resting state functional data were pre-processed and analysed using the FMRI Expert Analysis Tool (FEAT). The individual subject analysis involved: motion correction using MCFLIRT; slice-timing correction using Fourier space time-series phase-shifting; spatial smoothing using a Gaussian kernel of FWHM 6mm; grand-mean intensity normalisation of the entire 4D dataset by a single multiplicative factor; highpass temporal filtering (Gaussian-weighted least-squares straight line fitting, with sigma = 100 s); Gaussian lowpass temporal filtering, with sigma = 2.8s

##### Task based fMRI (Experiment 1)

First level analyses in experiment 1 modelled 6 explanatory variables (EV’s). EV’s 1 and 2 modelled time periods in which participants completed the 0 back task or 1-back condition. EV’s 3 – 6 modeled the four extracted principal components by assigning the beta weight of each extracted principal component to each thought at the trial level. Thought probes were modelled in a 6 second time window (the minimum time period between 2 thought probes) before the onset of each thought probe.

Individual participant data was first entered into a higher-level fixed-effect analysis to measure and average neural response to the 6 EV’s across all 8 functional runs. Following this, the fixed level effects were entered into a group analysis using a mixed-effects design (FLAME, http://www.fmrib.ox.ac.uk/fsl). Z stat maps were generated for each EV; 0 back task, 1-back task, PC1, PC2, PC3 and PC4. We also defined task specific neural responses by contrasting z stat maps for 0 back > 1-back and 1-back > 0 back conditions. These maps were then registered to a high resolution T1-anatomical image and then onto the standard MNI brain (ICBM152).

##### Functional Connectivity Analysis

*Region of Interest (ROI) Selection and Mask Creation* We selected seed regions of interest in the functional connectivity analysis based on the significant contrasts (cluster corrected at z>3.1) arising from the task based fMRI in experiment 1 (see figure 3 A and B): 0 back>1- back. To carry out the seed based analysis we binarised the Z>3.1 cluster corrected ROI masks and extracted the time series of these regions during the resting-state session. These time series were then used as explanatory variables in connectivity analyses at the single subject level. In these analyses, we entered 11 nuisance regressors; the top five principal components extracted from white matter (WM) and cerebrospinal fluid (CSF) masks based on the CompCor method [34], six head motion parameters and spatial smoothing (Gaussian) was applied at 6mm (FWHM). WM and CSF masks were generated from each individual’s structural image [35]. No global signal regression was performed, following the method implemented in [36].

We related ROI connectivity patterns to inter-individual variations in different types of thought using a multiple regression model, in which the connectivity maps was the dependent variable and z scores describing the four thought types as the explanatory variables: (i) PC1 ‘Detail’ 0 back, (ii) PC1 ‘Detail’ 1-back, (iii) PC2 ‘TUT’ 0 back, (iv) PC2 ‘TUT’ 1-back, (v) PC3 ‘Modality’ 0 back, (vi) PC3 ‘Modality’ 1-back, (vii) PC4 ‘Emotion’ 0 back, (viii) PC4 ‘Emotion’ 1-back. We included mean frame displacement [38] in our group level regressions to rule out spurious effects. These analyses were carried out using FMRIB’s Local Analysis of Mixed Effects (FLAME1).

#### Multiple comparison correction

In all fMRI analyses we used a cluster forming threshold of Z = 3.1 and controlled for family wise error at p<.05 [37].

##### Meta analytic decoding

We compared unthresholded functional connectivity activation profiles to those of previous studies using the Neurosynth decoder (http://www.neurosynth.org/decode/ see [10] for further details). To produce our word clouds we manually extracted the top ten task descriptions (based on frequency) for each unthresholded z map (we manually excluded the names of brain regions or MRI methods) to generate the word clouds in Figure 4.

## Author contributions

JS, EJ, HTW and MS designed the experiment. MS provided reactants and methods for the online sampling experiment. TK provided reactants and methods for the functional connectivity experiment. CM, MS, TK and HTW collected the data. MS analysed the data. All authors contributed to the writing of the paper.

## Acknowledgments

Thanks to the members of the Memory and Thought Lab for their help with the data collection, and to Michael Mrazek for providing the measure of Ravens Matrices. EJ was supported by BBSRC (BB/J006963/1) and the ERC (SEMBIND - 283530). JS was supported by the ERC (WANDERINGMINDS - 646927) and the Volkswagen Foundation (Wandering Minds - 89440 and 89439) and a grant from the John Templeton Foundation, “Prospective Psychology Stage 2: A Research Competition”. TK and CM were supported by a Departmental Studentship from the Psychology Department.

## Supplementary Materials

**ST1.** Correlations between the decomposition solutions from lab and scanner. These correlations were computed using the rotated component matrix scores for each principal component independently.

|  | Detail | TUT | Modality | Emotion |
| --- | --- | --- | --- | --- |
| r | 0.74 | 0.96 | 0.73 | 0.64 |
| p | <0.005 | <.001 | 0.0046 | 0.018 |

**ST2.** Multiple Dimension Experience Sampling questions in 0-back / 1-back task in both Experiments.

| Dimensions | Questions | 0 | 1 |
| --- | --- | --- | --- |
| Focus | My thoughts were focused on the task I was performing. | Not at all | Completely |
| Future | My thoughts involved future events. | Not at all | Completely |
| Past | My thoughts involved past events. | Not at all | Completely |
| Self | My thoughts involved myself. | Not at all | Completely |
| Other | My thoughts involved other people. | Not at all | Completely |
| Emotion | The content of my thoughts was: | Negative | Positive |
| Images | My thoughts were in the form of images. | Not at all | Completely |
| Words | My thoughts were in the form of words. | Not at all | Completely |
| Vivid | My thoughts were vivid as if I was there. | Not at all | Completely |
| Vague | My thoughts were detailed and specific. | Not at all | Completely |
| Habit | This thought has recurrent themes similar to those I have had before. | Not at all | Completely |
| Evolving | My thoughts tended to evolve in a series of steps. | Not at all | Completely |
| Spontaneous | My thoughts were: | Spontaneous | Deliberate |

**SF1.**
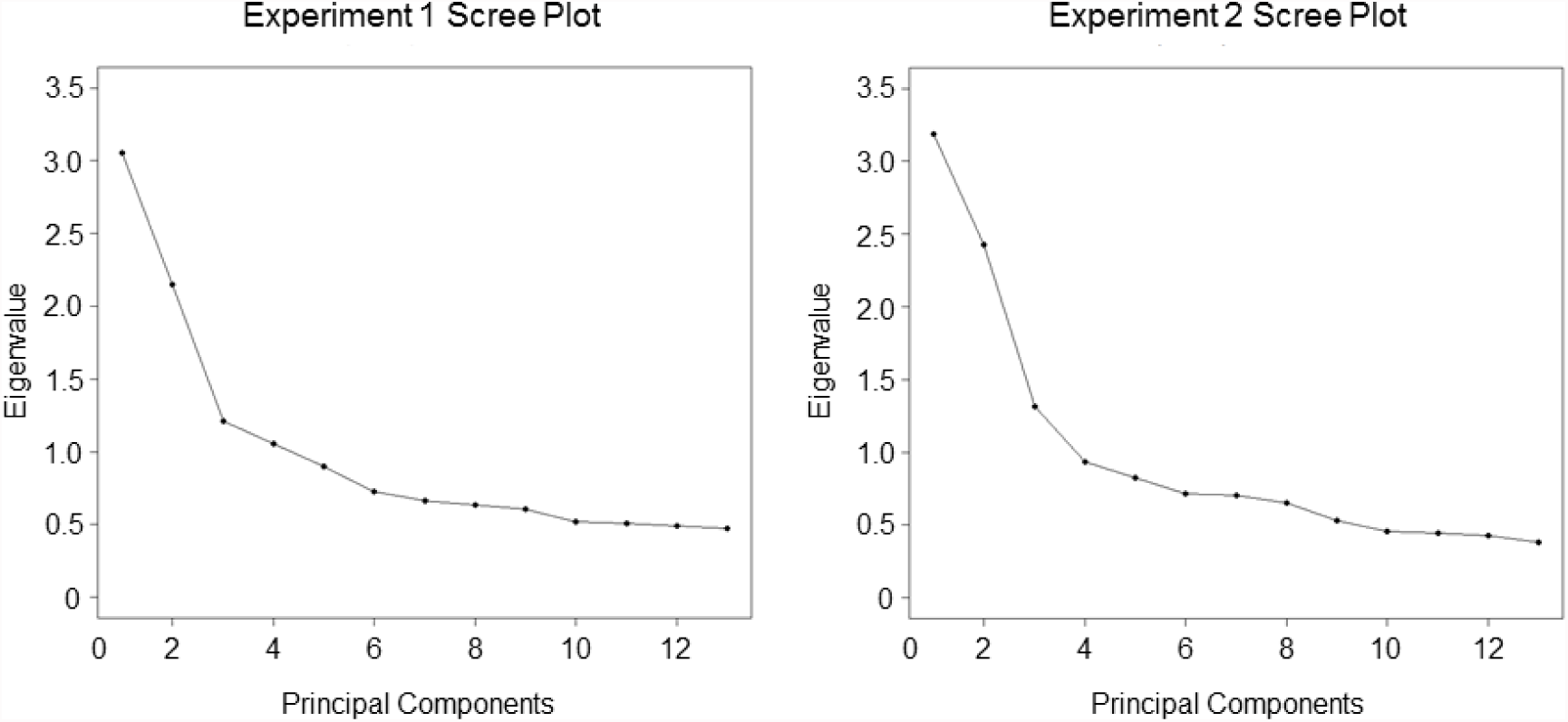
Scree plots displaying Eigenvalue scores for Principal components extracted using varimax rotation in Experiments 1 and 2

**SF2.**
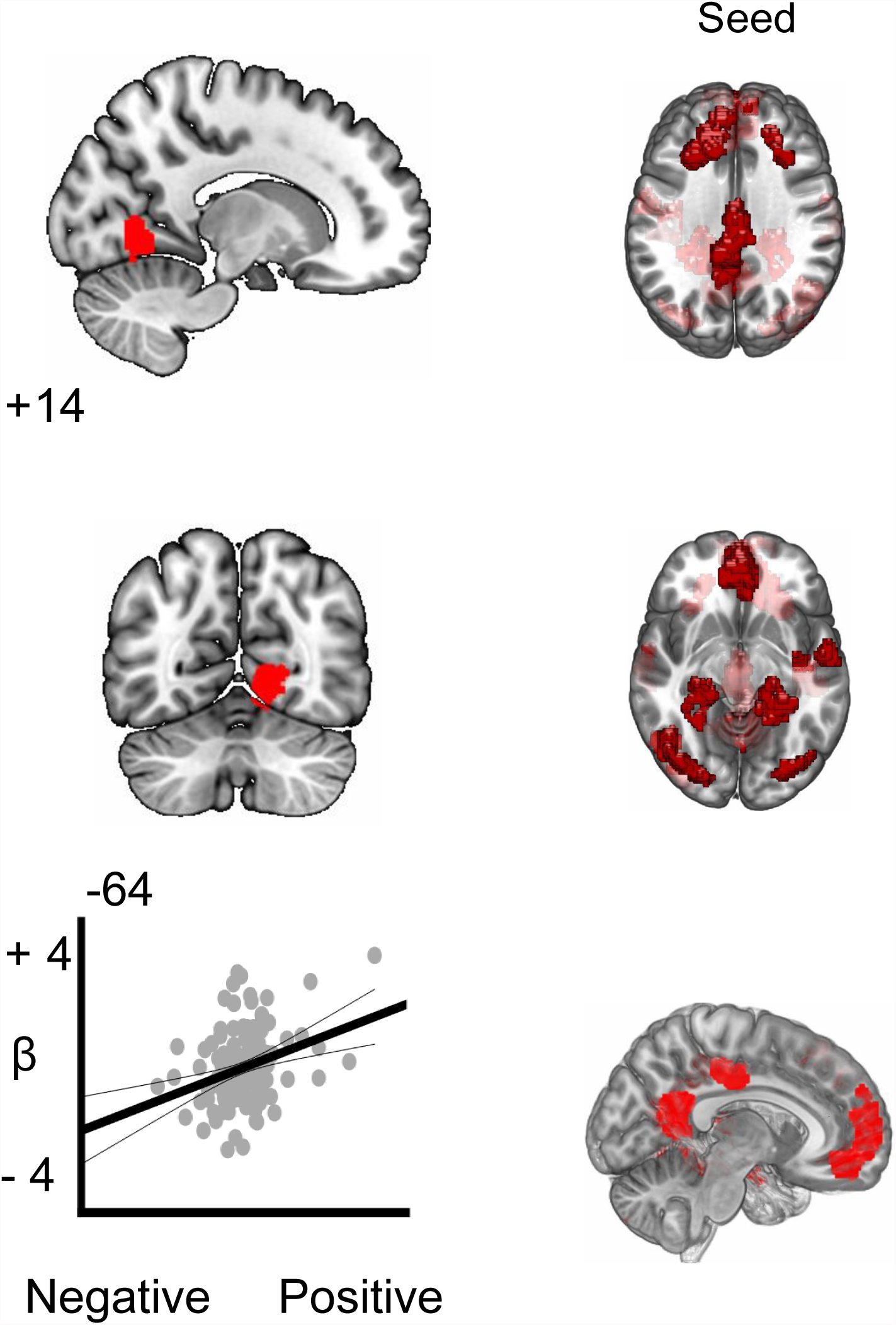
Results of group level functional connectivity analysis showing the positive correlation between activity in the lingual gyrus and regions showing enhanced activity in the non demanding task conditions from Experiment one. Brain images were thresholded at Z = 3.1 and multiple comparisons were controlled at p<.05.

**SF3.**
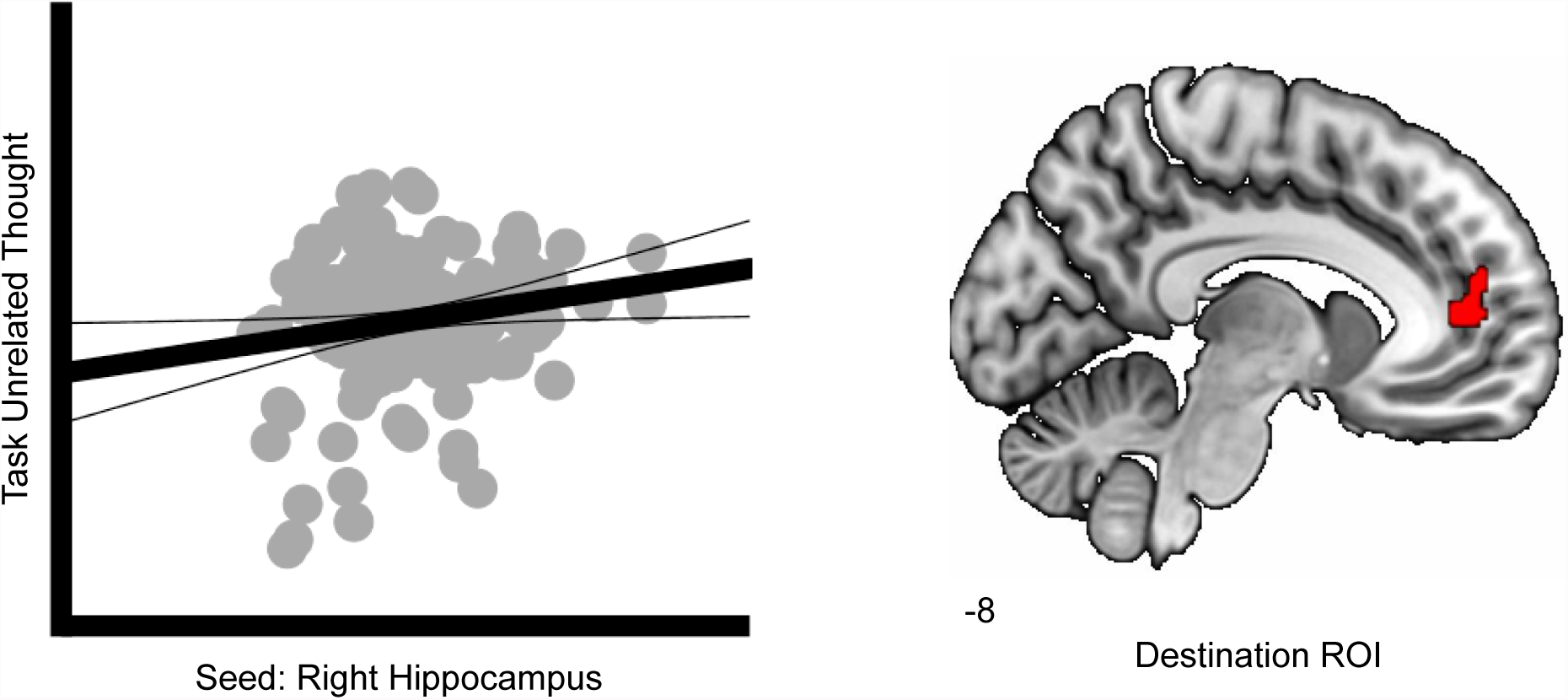
Correlation replicating the relationship between connectivity of the right hippocampus and the region of ventral mPFC identified by Karapanagiotidis and colleagues (r = .173, p<.05). This ventral region of mPFC showed greater connectivity to a hippocampus seed region for episodic aspects of mind-wandering.

**SF4.**
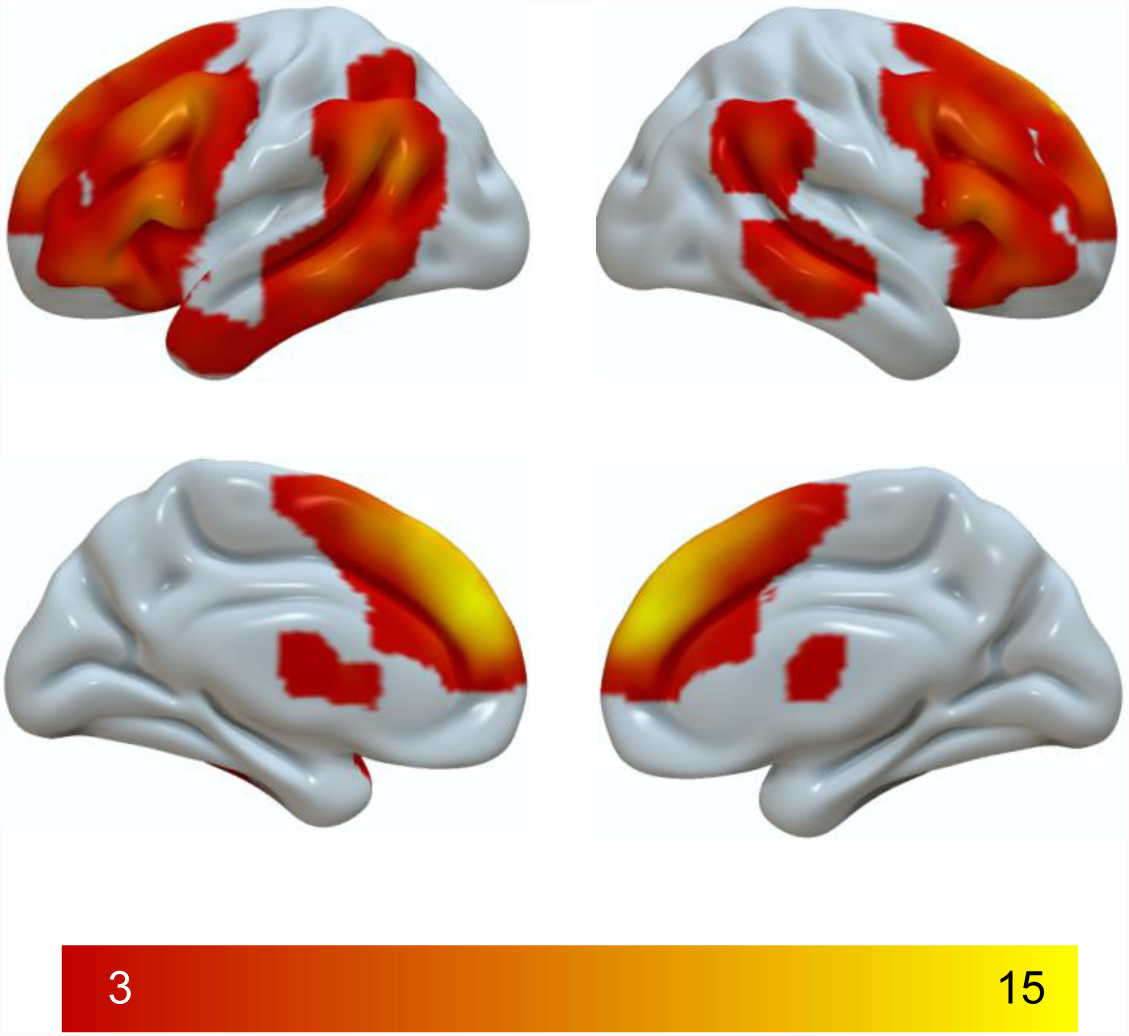
Cluster corrected map regions showing significantly stronger correlations with dorsal than ventral mPFC. The spatial map is thresholded at z = 3.1 and corrected for multiple comparisons at p<.05.

## References

1. Killingsworth, M.A. and D.T. Gilbert, A wandering mind is an unhappy mind. Science, 2010. 330(6006): p. 932.

2. Poerio, G.L., P. Totterdell, and E. Miles, Mind-wandering and negative mood: does one thing really lead to another? Conscious Cogn, 2013. 22(4): p. 1412-21.

3. Kane, M.J., et al., For whom the mind wanders, and when: an experience-sampling study of working memory and executive control in daily life. Psychol Sci, 2007. 18(7): p. 614-21.

4. Baird, B., et al., Inspired by distraction: mind wandering facilitates creative incubation. Psychol Sci, 2012. 23(10): p. 1117-22.

5. Baird, B., J. Smallwood, and J.W. Schooler, Back to the future: autobiographical planning and the functionality of mind-wandering. Conscious Cogn, 2011. 20(4): p. 1604-11.

6. Engert, V., J. Smallwood, and T. Singer, Mind your thoughts: associations between self-generated thoughts and stress-induced and baseline levels of cortisol and alpha-amylase. Biol Psychol, 2014. 103: p. 283-91.

7. McVay, J.C. and M.J. Kane, Conducting the train of thought: working memory capacity, goal neglect, and mind wandering in an executive-control task. J Exp Psychol Learn Mem Cogn, 2009. 35(1): p. 196-204.

8. Smallwood, J. and J.W. Schooler, The science of mind wandering: empirically navigating the stream of consciousness. Annu Rev Psychol, 2015. 66: p. 487-518.

9. Christoff, K., et al., Mind-wandering as spontaneous thought: a dynamic framework. Nat Rev Neurosci, 2016. 17(11): p. 718-731.

10. Smallwood, J., Distinguishing how from why the mind wanders: a process-occurrence framework for self-generated mental activity. Psychol Bull, 2013. 139(3): p. 519-35.

11. Mrazek, M.D., et al., The role of mind-wandering in measurements of general aptitude. J Exp Psychol Gen, 2012. 141(4): p. 788-98.

12. McVay, J.C. and M.J. Kane, Does mind wandering reflect executive function or executive failure? Comment on Smallwood and Schooler (2006) and Watkins (2008). Psychol Bull, 2010. 136(2): p. 188-97; discussion 198-207.

13. Levinson, D.B., J. Smallwood, and R.J. Davidson, The persistence of thought: evidence for a role of working memory in the maintenance of task-unrelated thinking. Psychol Sci, 2012. 23(4): p. 375-80.

14. Sanders, J.G., et al., Can I get me out of my head? Exploring strategies for controlling the self-referential aspects of the mind-wandering state during reading. Q J Exp Psychol (Hove), 2017. 70(6): p. 1053-1062.

15. Mrazek, M.D., et al., Mindfulness training improves working memory capacity and GRE performance while reducing mind wandering. Psychol Sci, 2013. 24(5): p. 776-81.

16. Rahl, H.A., et al., Brief mindfulness meditation training reduces mind wandering: The critical role of acceptance. Emotion, 2017. 17(2): p. 224-230.

17. Redshaw, J. and T. Suddendorf, Foresight beyond the very next event: four-year-olds can link past and deferred future episodes. Front Psychol, 2013. 4: p. 404.

18. Vygotsky, L.S., Imagination and creativity in childhood. Journal of Russian & East European Psychology, 2004. 42(1): p. 7-97.

19. Suddendorf, T. and M.C. Corballis, Behavioural evidence for mental time travel in nonhuman animals. Behav Brain Res, 2010. 215(2): p. 292-8.

20. Suddendorf, T. and M.C. Corballis, Mental time travel and the evolution of the human mind. Genet Soc Gen Psychol Monogr, 1997. 123(2): p. 133-67.

21. Suddendorf, T. and M.C. Corballis, The evolution of foresight: What is mental time travel, and is it unique to humans? Behav Brain Sci, 2007. 30(3): p. 299-313; discussion 313-51.

22. Bastian, M., et al., Language facilitates introspection: Verbal mind-wandering has privileged access to consciousness. Conscious Cogn, 2017. 49: p. 86-97.

23. Stawarczyk, D., H. Cassol, and A. D’Argembeau, Phenomenology of future-oriented mind-wandering episodes. Front Psychol, 2013. 4: p. 425.

24. Andrews-Hanna, J.R., J. Smallwood, and R.N. Spreng, The default network and self-generated thought: component processes, dynamic control, and clinical relevance. Ann N Y Acad Sci, 2014. 1316: p. 29-52.

25. Fox, K.C., et al., The wandering brain: Meta-analysis of functional neuroimaging studies of mind-wandering and related spontaneous thought processes. NeuroImage, 2015. 111: p. 611-621.

26. Stawarczyk, D. and A. D’Argembeau, Neural correlates of personal goal processing during episodic future thinking and mind-wandering: An ALE meta-analysis. Human brain mapping, 2015.

27. Raichle, M.E., et al., A default mode of brain function. Proceedings of the National Academy of Sciences of the United States of America, 2001. 98(2): p. 676-82.

28. Perogamvros, L., et al., The Phenomenal Contents and Neural Correlates of Spontaneous Thoughts across Wakefulness, NREM Sleep, and REM Sleep. J Cogn Neurosci, 2017: p. 1-12.

29. Raij, T.T. and T.J.J. Riekki, Dorsomedial prefontal cortex supports spontaneous thinking per se. Hum Brain Mapp, 2017. 38(6): p. 3277-3288.

30. Tusche, A., et al., Classifying the wandering mind: revealing the affective content of thoughts during task-free rest periods. Neuroimage, 2014. 97: p. 107-116.

31. Stawarczyk, D., et al., Neural correlates of ongoing conscious experience: both task-unrelatedness and stimulus-independence are related to default network activity. PLoS One, 2011. 6(2): p. e16997.

32. Christoff, K., et al., Experience sampling during fMRI reveals default network and executive system contributions to mind wandering. Proceedings of the National Academy of Sciences, 2009. 106(21): p. 8719-8724.

33. Addis, D.R. and D.L. Schacter, Constructive episodic simulation: Temporal distance and detail of past and future events modulate hippocampal engagement. Hippocampus, 2008. 18(2): p. 227-237.

34. Addis, D.R., A.T. Wong, and D.L. Schacter, Remembering the past and imagining the future: Common and distinct neural substrates during event construction and elaboration. Neuropsychologia, 2007. 45(7): p. 1363-1377.

35. Smallwood, J., L. Nind, and R.C. O’Connor, When is your head at? An exploration of the factors associated with the temporal focus of the wandering mind. Consciousness and cognition, 2009. 18(1): p. 118-25.

36. Konishi, M., et al., Shaped by the Past: The Default Mode Network Supports Cognition that Is Independent of Immediate Perceptual Input. PLoS One, 2015. 10(6): p. e0132209.

37. Karapanagiotidis, T., et al., Tracking thoughts: Exploring the neural architecture of mental time travel during mind-wandering. Neuroimage, 2017. 147: p. 272-281.

38. Smallwood, J., et al., Representing Representation: Integration between the Temporal Lobe and the Posterior Cingulate Influences the Content and Form of Spontaneous Thought. PLoS One, 2016. 11(4): p. e0152272.

39. Christoff, K., et al., Experience sampling during fMRI reveals default network and executive system contributions to mind wandering. Proc Natl Acad Sci U S A, 2009. 106(21): p. 8719-24.

40. Yeo, B.T., et al., The organization of the human cerebral cortex estimated by intrinsic functional connectivity. Journal of neurophysiology, 2011. 106(3): p. 1125-1165.

41. Seeley, W.W., et al., Dissociable intrinsic connectivity networks for salience processing and executive control. J Neurosci, 2007. 27(9): p. 2349-56.

42. Duncan, J., The multiple-demand (MD) system of the primate brain: mental programs for intelligent behaviour. Trends Cogn Sci, 2010. 14(4): p. 172-9.

43. Crittenden, B.M. and J. Duncan, Task difficulty manipulation reveals multiple demand activity but no frontal lobe hierarchy. Cereb Cortex, 2014. 24(2): p. 532-40.

44. Andrews-Hanna, J.R., et al., Functional-anatomic fractionation of the brain’s default network. Neuron, 2010. 65(4): p. 550-62.

45. Yarkoni, T., et al., Large-scale automated synthesis of human functional neuroimaging data. Nat Methods, 2011. 8(8): p. 665-70.

46. Pulvermuller, F., Brain mechanisms linking language and action. Nat Rev Neurosci, 2005. 6(7): p. 576-82.

47. Pulvermuller, F., et al., Functional links between motor and language systems. Eur J Neurosci, 2005. 21(3): p. 793-7.

48. Gallese, V. and G. Lakoff, The Brain’s concepts: the role of the Sensory-motor system in conceptual knowledge. Cogn Neuropsychol, 2005. 22(3): p. 455-79.

49. Beisteiner, R., et al., Mental representations of movements. Brain potentials associated with imagination of hand movements. Electroencephalogr Clin Neurophysiol, 1995. 96(2): p. 183-93.

50. Medea, B., et al., How do we decide what to do? Resting-state connectivity patterns and components of self-generated thought linked to the development of more concrete personal goals. Exp Brain Res, 2016.

51. Alderson-Day, B. and C. Fernyhough, Inner Speech: Development, Cognitive Functions, Phenomenology, and Neurobiology. Psychol Bull, 2015. 141(5): p. 931-65.

52. Vallar, G. and A.D. Baddeley, Fractionation of working memory: Neuropsychological evidence for a phonological short-term store. Journal of Verbal Learning and Verbal Behavior, 1984. 23(2): p. 151-161.

53. Ralph, M.A., et al., The neural and computational bases of semantic cognition. Nat Rev Neurosci, 2017. 18(1): p. 42-55.

54. Binder, J.R., et al., Where is the semantic system? A critical review and meta-analysis of 120 functional neuroimaging studies. Cereb Cortex, 2009. 19(12): p. 2767-96.

55. Smallwood, J., F.J. Ruby, and T. Singer, Letting go of the present: mind-wandering is associated with reduced delay discounting. Conscious Cogn, 2013. 22(1): p. 1-7.

56. Bernhardt, B.C., et al., Medial prefrontal and anterior cingulate cortical thickness predicts shared individual differences in self-generated thought and temporal discounting. Neuroimage, 2014. 90: p. 290-7.

57. Rummel, J. and C.D. Boywitt, Controlling the stream of thought: working memory capacity predicts adjustment of mind-wandering to situational demands. Psychon Bull Rev, 2014. 21(5): p. 1309-15.

58. Smallwood, J., M. McSpadden, and J.W. Schooler, When attention matters: the curious incident of the wandering mind. Mem Cognit, 2008. 36(6): p. 1144-50.

59. Ottaviani, C., et al., Worry as an adaptive avoidance strategy in healthy controls but not in pathological worriers. Int J Psychophysiol, 2014. 93(3): p. 349-55.

60. Sormaz, M., et al., Knowing what from where: Hippocampal connectivity with temporoparietal cortex at rest is linked to individual differences in semantic and topographic memory. Neuroimage, 2017. 152: p. 400-410.

